# Cell-free mutant analysis combined with structure prediction of a lasso peptide biosynthetic protein B2

**DOI:** 10.1101/2022.04.06.487251

**Authors:** Almasul Alfi, Aleksandr Popov, Ashutosh Kumar, Kam Y. J. Zhang, Svetlana Dubiley, Konstantin Severinov, Shunsuke Tagami

## Abstract

Biochemical and structural analyses of purified proteins are essential for the understanding of their properties. However, many proteins are unstable and difficult to purify, hindering their characterization. The B2 proteins of the lasso peptide biosynthetic pathways are cysteine proteases that cleave precursor peptides during the maturation process. The B2 proteins are poorly soluble and no experimentally-solved structures are available. Here, we performed a rabid semi-comprehensive mutational analysis of the B2 protein from the thermophilic actinobacterium, *Thermobifida fusca* (TfuB2) using a cell-free transcription/translation system, and compared the results with the structure prediction by AlphaFold2. Analysis of 34 TfuB2 mutants with substitutions of hydrophobic residues confirmed the accuracy of the predicted structure, and revealed a hydrophobic patch on the protein surface, which likely serves as the binding site of the partner protein, TfuB1. Our results suggest that the combination of rapid cell-free mutant analyses with precise structure predictions can greatly accelerate structure-function research of proteins for which no structures are available.

---

Lasso peptides are a class of ribosomally synthesized and post-translationally modified peptides (RiPPs) with a unique 3D interlocked structure (Figure 1A).^1–3^ The N-terminal amido group of a lasso peptide is conjugated with the side chain of an acidic residue (Asp/Glu) at positions 7– 10 by an isopeptide bond to form a ring structure; the C-terminal linear part is threaded through the ring and is topologically trapped in the position. Lasso peptide biosynthesis is performed by proteins encoded in a gene cluster that also includes genes encoding the linear peptide precursor and the export machinery components.^1–4^ The precursor peptide (the A peptide) consists of the N-terminal leader part (A-Leader) and the C-terminal core part (A-Core). First, the leader part is recognized and cleaved by the B1 and B2 proteins, which are sometimes fused into a single protein, B. The C protein then catalyzes the formation of the N-terminal ring in the core. Although the detailed mechanism of leader recognition was revealed by the structures of the B1 protein complexed with the leader peptide,^5,6^ the mechanism of the subsequent lasso formation processes has remained elusive.

**Figure 1.**
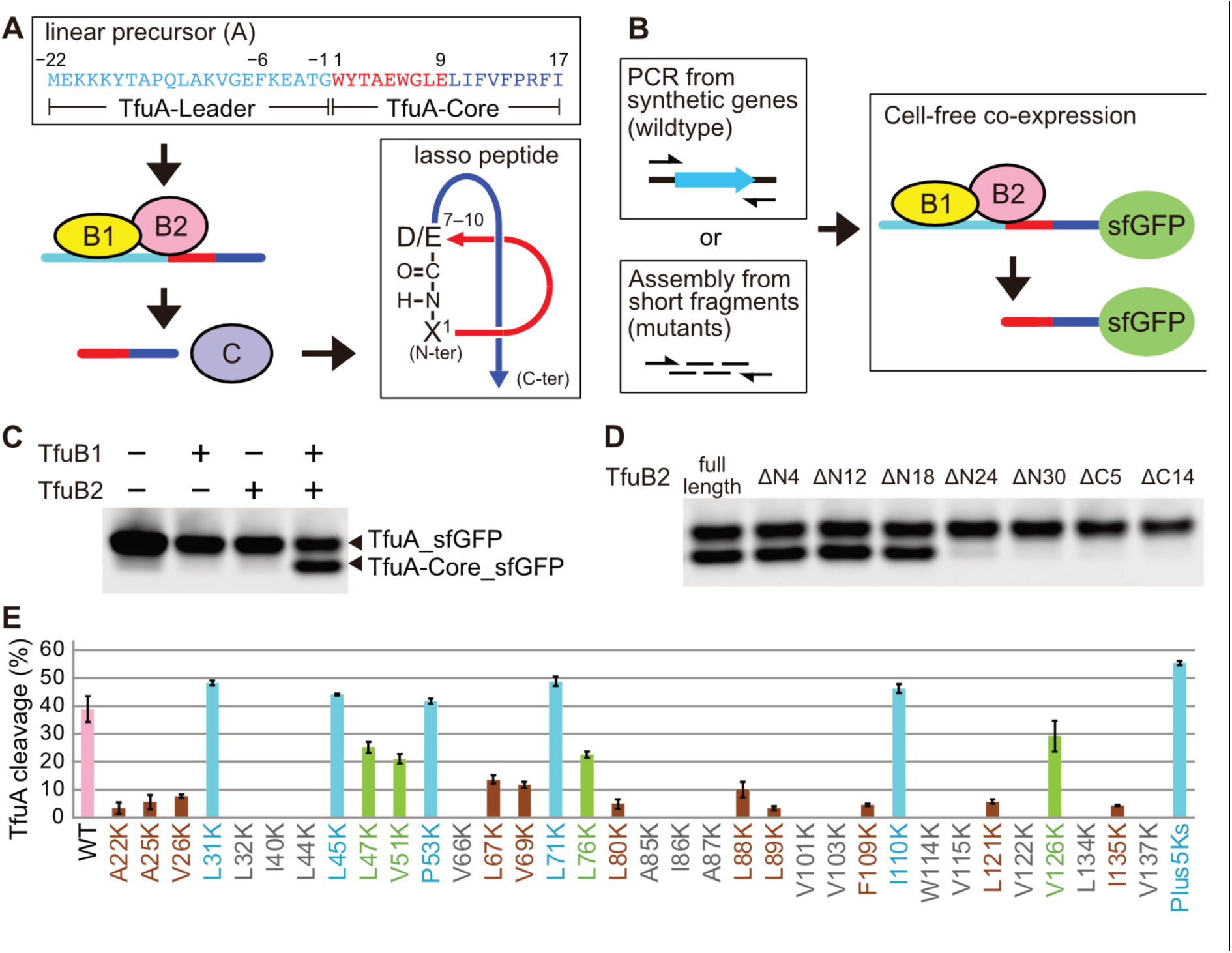
Cell-free mutant analyses of the B2 protein. Schematic depictions of (A) the biosynthetic pathway of lasso peptide and (B) the cell-free mutant analysis of the B2 protein. (C) Cell-free cleavage of TfuA_sfGFP in the presence of co-expressed TfuB1 and TfuB2. (D) Cleavage of TfuA_sfGFP by the truncation variants of TfuB2 (full length: 1– 138; ΔN4: 5–138; ΔN12: 13–138; ΔN18: 19–138; ΔN24: 25–138; ΔN30: 31–138; ΔC5: 1– 133; ΔC14: 1–124). (E) Average cleavage of TfuA_sfGFP by the indicated mutants in the PURE system. Mutants are colored according to their average cleavage activities (cyan: > 40%; lime: >20%; brown: <15%, gray: no activity detected). Error bars represent S.D. (N ≧ 3).

Although the B2 and C proteins from several different bacteria are reportedly active in heterologous *in vivo* systems,^7^ their structural analyses have been hindered by their insolubility and instability. Fusions with large tags increasing solubility are usually required for the purification of the B2 and C proteins, such as McjB and McjC from *Escherichia coli* and TfuB2 and TfuC from *Thermobifida fusca*.^8,9^ The TfuB2•TfuC complex was reportedly purified without solubility tags, but still eluted in the void volume from a size exclusion column,^10^ indicating that it forms macromolecular aggregates. At the time of this writing, none of the B2 and C proteins have been purified in sufficient qualities and quantities for crystallization.

One possible way to bypass this problem and accelerate research on the enigmatic process of lasso peptide maturation would be to use the recently developed programs for precise structure prediction, i.e., AlphaFold2 and RoseTTAFold.^11,12^ However, there are still many cases where these programs fail to predict the correct structures. Further, predictions must be confirmed by experiments that should ideally be as speedy as the structure prediction.

In standard mutagenesis experiments, one must construct expression plasmids containing cloned genes of interest, introduce and validate site-specific mutations, transform them into a production host, and purify the mutant proteins for functional analysis. These time-consuming procedures can be omitted by polymerase cycling assembly (also known as assembly PCR) of mutant genes from short DNA fragments,^13^ followed by protein expression in a cell-free *in vitro* transcription-translation system (IVTT).^14^ IVTT systems can also be used to monitor the activities of enzymes without purification.^15–17^ For example, the IVTT system reconstituted from purified components (the PURE system)^18^ was used to monitor the activities of ribosomes containing incomplete sets of ribosomal proteins, by measuring the fluorescence intensity of superfolder green fluorescent protein (sfGFP) produced by the ribosome variants.^19^ If these technologies (*in vitro* mutagenesis, IVTT, and activity measurement without purification) are combined, then the analysis of multiple mutant proteins could, in theory, be finished in just one day.^20^

In this report, we performed structure prediction by AlphaFold2 and semi-comprehensive cell-free mutant analysis to achieve quick and reliable characterization of the structure of TfuB2. The results of mutagenesis and structure prediction were highly consistent with each other, confirming the accuracy of the predicted model. Analysis of the validated structural model allowed us to identify the likely binding site for the B1 protein on the TfuB2 surface. These observations were further confirmed for the B2 protein from a firmicute, *Bacillus pseudomycoides* (PsmB2), demonstrating that the structural features of the B2 protein are conserved in a wide variety of organisms.

## RESULTS AND DISCUSSION

### Quick functional analysis of TfuB2 mutants using a cell-free translation system

We set to develop a cell-free PURE system-based pipeline for quick analyses of the B2 protein mutants (Figure 1B), with the original goal of engineering more soluble variants for structural analysis. We chose the lasso peptide biosynthetic system from *T. fusca*, as the structure of the B1 protein (TfuB1) was successfully determined^5^ and *in vitro* synthesis of the lasso peptide (fusilassin/fuscanodin) in the PURE system was reported.^21^ Synthetic genes of TfuB1, TfuB2, and the precursor peptide fused with sfGFP (TfuA_sfGFP) were amplified by PCR using primers containing a T7 promoter and an appropriately positioned ribosome binding site (Supplementary Tables 1–3). Proteins encoded by resulting DNA molecules were co-expressed in the PURE system at 37 ºC for 1 hour and then incubated at 50 ºC, the temperature for the TfuB2 activity test. The fluorescent band of TfuA_sfGFP cleaved by TfuB2 (TfuA-Core_sfGFP) was observed only in the reactions where both TfuB1 and TfuB2 were co-expressed (Figure 1C).

Constructs encoding truncated versions of TfuB2 were next tested in the same system to identify a minimal fragment capable of the peptidase activity (Figure 1D). The N-terminal truncations extending up to residue Thr18 (TfuB2-ΔN4, TfuB2-ΔN12, and TfuB2-ΔN18) were almost as active as the full-length protein. More extensive N-terminal truncations (TfuB2-ΔN24 and TfuB2-ΔN30) showed diminished peptidase activities, while the two C-terminal truncations tested (TfuB2-ΔC5, TfuB2-ΔC14) were inactive. Thus, at the level of resolution of our analysis, the essential part of TfuB2 includes residues 19–138.

Next, genes encoding 34 randomly-chosen substitutions of TfuB2 hydrophobic residues for lysine were assembled (Figure 1B, Supplementary Tables 1–3). These mutations were introduced in the TfuB2-ΔN4 background to reduce the number of required DNA fragments. The amounts of the mutant proteins produced in the PURE system were not very different from each other or the “wild-type” TfuB2-ΔN4 protein (Supplementary Figure 1). Roughly 40% of the precursor peptide was cleaved by the parental TfuB2-ΔN4 at our condition (Figure 1E). Five mutants (L31K, L45K, P53K, L71K, I110K) showed activities equivalent to or slightly higher than that of the wild-type TfuB2. Four mutants (L47K, V51K, L76K, V126K) showed reduced but significant activities (20–30% cleavage of the TfuA-sfGFP substrate). Eleven mutants (A22K, A25K, V26K, L67K, V69K, L80K, L88K, L89K, F109K, L121K, and I135K) showed strongly reduced activities (<15% cleavage of the substrate). Finally, fourteen mutants (L32K, I40K, L44K, V66K, A85K, I86K, A87K, V101K, V103K, W114K, V115K, V122K, L134K, and V137K) completely lost the activity, indicating that the substituted amino acid residues are essential for the protein structure or interactions with other molecules (TfuA and TfuB1).

We combined the five substitutions that did not decrease the peptidase activity (L31K, L45K, P53K, L71K, I110K) into a single variant (TfuB2-ΔN4 Plus5Ks) and confirmed that its activity was improved compared to the wild-type TfuB2 (Figure 1E, ∼55% cleavage). We expected that TfuB2 Plus5Ks would also have a higher solubility than the wild-type TfuB2. Unfortunately, and similarly to the wild-type, TfuB2-ΔN4 Plus5Ks was completely insoluble (Supplementary Figure 2) and was thus not suitable for structural analysis.

### Comparison between the mutant analysis and structure prediction of TfuB2

After we failed to engineer a soluble TfuB2 variant, AlphaFold2 was released and became available^11^. We thereby predicted the structure of TfuB2 by the program and compared the prediction with the results of the mutational analysis described above (Figure 2). TfuB2 is predicted to be a single-domain protein composed of an N-terminal loop (1–16), an α-helical part (17–96), and a C-terminal β-sheet (97–138) (Figure 2A). The confidence of the prediction reported by the program was very high except for the N-terminal loop (Supplementary Figure 3). The predicted local-distance difference test score (pLDDT score) was found to be more than 90 for all five predicted models. The N-terminal region of TfuB2 found to be non-essential for the enzymatic activity (residues 1–18, Figure 1D) almost corresponds to the predicted N-terminal loop, indicating that it is flexible and not required for folding of the rest of the protein. The nine mutations that either had no effect or retained significant (>20% cleavage) activities – L31K, L45K, L47K, V51K, P53K, L71K, L76K, I110K, V126K – all mapped on the surface of the predicted structure (Figure 2B). In contrast, 18 of the 25 mutations with no or strongly reduced (<15% cleavage) activities – A22K, A25K, V26K, L32K, I40K, L44K, V66K, L67K, V69K, L80K, A85K, I86K, A87K, L88K, L89K, V115K, V122K, and V137K – are in the hydrophobic core of the protein (Figure 2C). The high consistency between the predicted structure and the semi-comprehensive mutagenesis confirms the accuracy of the structure prediction.

**Figure 2.**
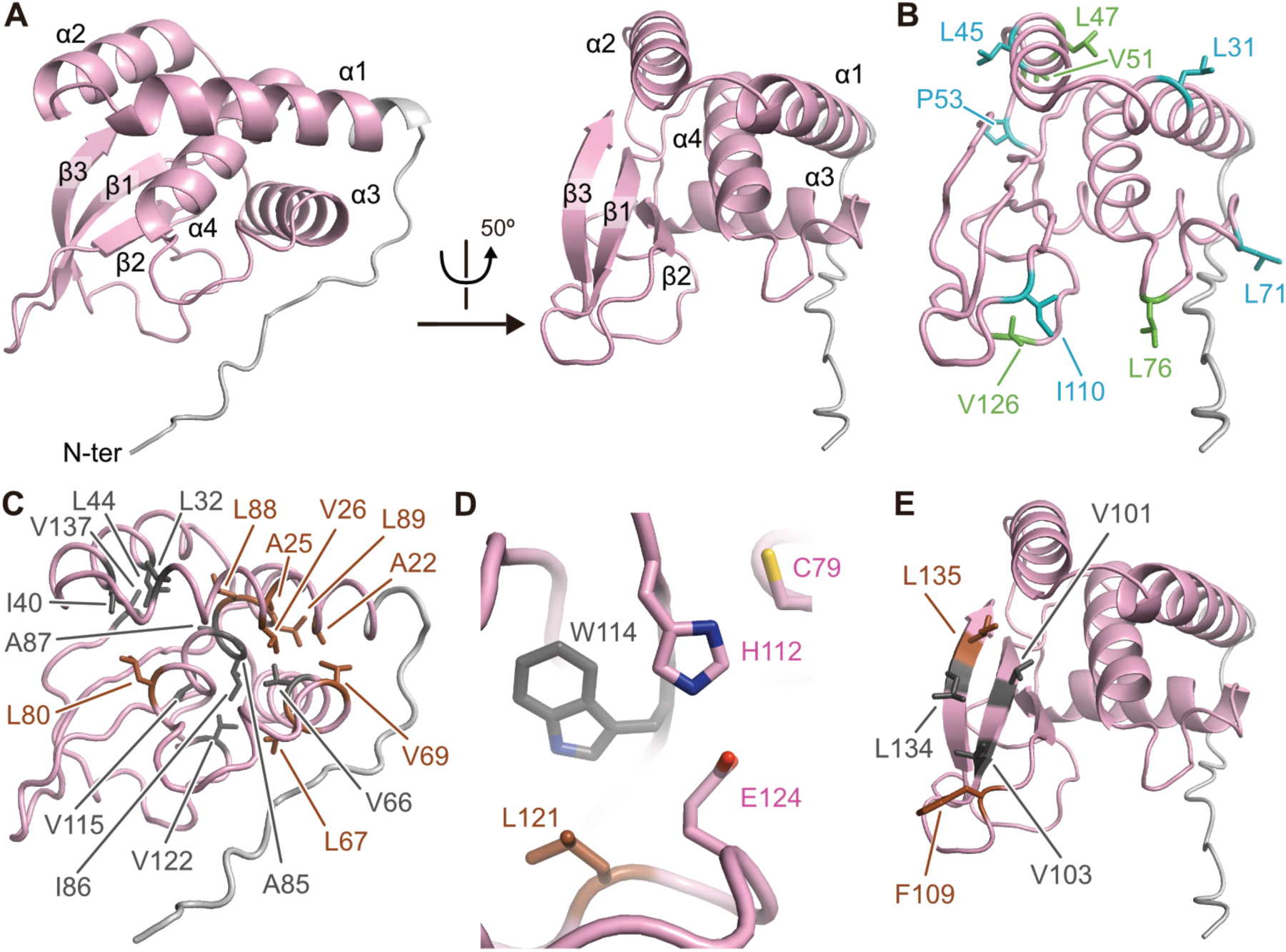
The predicted structure of TfuB2. (A) Overall structure of TfuB2 predicted by AlphaFold2. The functionally dispensable N-terminal part is colored light gray. (B) Residues whose substitutions preserved activity (>20% TfuA-sfGFP cleavage) are mapped on the predicted TfuB2 structure. (C) Positions of residues whose substitutions strongly reduced activity (<15% TfuA-sfGFP cleavage) and which are located in the hydrophobic cores are mapped on the predicted TfuB2 structure. (D) Close-up view of the active site. (E) Residues whose substitutions abolished activity (<5% TfuA-sfGFP cleavage) and that are clustered on the protein surface are mapped on the TfuB2 structural model. The mutated residues are colored in the same way as in Figure 1E.

Two other lysine mutations with no or barely detectable activity (W114K and L121K) are located around the predicted active-site residues (Figure 2D). The B2 proteins are members of transglutaminase-like peptidases and have a conserved Cys-His-Asp/Glu triad at the active center, a typical feature of the peptidase family.^4,8,22–24^ The active-site triad is formed by Cys79, His112, and Glu124 in the predicted structure. It was previously reported that alanine substitutions of Cys79 and His112 completely inactivated the B2 protein, while the E124A substitution was active.^9^ We also tested these alanine substitutions in our cell-free analysis (Supplementary Figure 4). The C79A and H112A mutants had no activity, while E124A showed significantly reduced but clearly detectable cleavage of TfuA (15.0% cleavage, S.D. = 0.3%, N = 3). In the predicted TfuB2 structure, Trp114 and Leu121 are contacting His112 and Glu124, respectively, and likely stabilizing the positions of these residues and the conformation of the loop between β2 and β3 (117–130) (Figure 2D and Supplementary Figure 5). It is thus likely that the absence of activity in W114K and L121K mutants is due to repositioning of the catalytic residues and distortion of the loop around the active site.

Interestingly, in the predicted structure of TfuB2, the other five hydrophobic residues whose lysine substitutions abolished the peptidase activity (V101, V103, F109, L134, and I135) form a surface-exposed hydrophobic patch (Figure 2E). It is highly likely that these residues are involved in interactions with TfuA and/or TfuB1. To get further insights into the function of this patch, we predicted the structure of the TfuA-Leader•TfuB1•TfuB2 complex by using AlphaFold2 (“model_preset = multimer” option) (Figure 3 and Supplementary Figure 6).^11,25^ The predicted structure must be at least partially correct, as the TfuA-Leader•TfuB1 moiety of the prediction is quite consistent with the previously solved crystal structure of the TfuA-Leader•TfuB1 complex (Supplementary Figure 7).^5^ In the predicted complex structure, the hydrophobic patch of TfuB2 forms an interaction surface with the intermolecular β-sheet jointly formed by TfuA-Leader•TfuB1 (Figure 3A, B). The intermolecular β-sheet was previously considered to be the interaction surface with TfuB2, as its hydrophobic residues (TfuA Phe−6, TfuB1 Val24, Lue26, Tyr33) are well conserved but not essential for the B1 protein to recognize TfuA-Leader.^5^ Mutations of some residues on the intermolecular β-sheet (TfuB1 L26A, G31A, Y33A) reportedly abolished the leader peptide cleavage by the B2 protein, without affecting the recognition of the leader peptide by TfuB1.^9^ Furthermore, the C-terminus of the TfuA-Leader, the site cleaved by TfuB2, is near the catalytic cysteine residue (TfuB2 Cys79, Figure 3A). Thus, all data obtained in the previous reports and our current work are consistent with the TfuA-Leader•TfuB1•TfuB2 complex model predicted by AlphaFold2.

**Figure 3.**
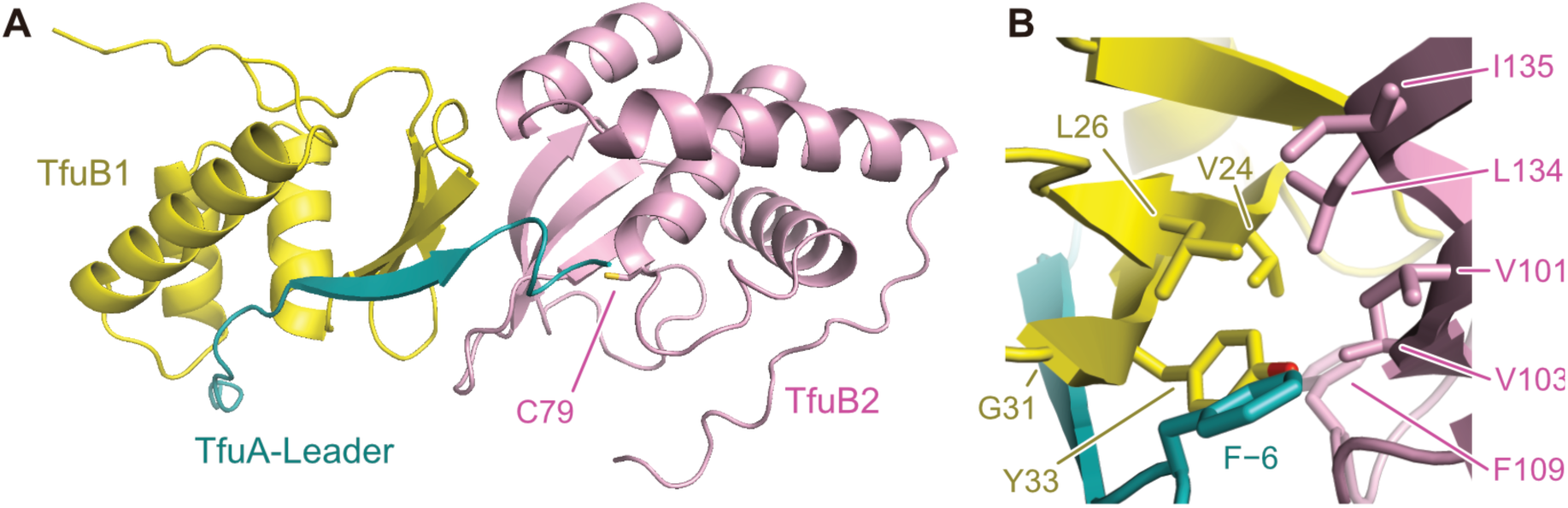
The predicted structure of the TfuA•TfuB1•TfuB2 complex. (A) The structural model of the TfuA•TfuB1•TfuB2 complex predicted by AlphaFold2. (B) A close-up view of the interaction surface between TfuA-Leader•TfuB1 and TfuB2.

### Structure prediction and mutant analysis of the B2 protein from a firmicute

To establish whether the predicted structural features can be generalized to other B2 proteins, we performed structure prediction and mutant analysis of the B2 protein from a firmicute, *Bacillus pseudomycoides* (PsmB2).^26^ The structure of PsmB2 predicted by AlphaFold2 had the same topology as TfuB2, except for the N-terminal PsmB2 residues 1–14, which were modeled as an α-helix with relatively low confidence (Figure 4A and Supplementary Figure 8). Because we could not observe the activity of PsmB2 in the cell-free system, we used the previously reported *in vivo* activity test to follow the functions of PsmB2 mutants (Figure 4B). In this experiment, the precursor peptide was conjugated with MBP and Trx (MBP_PsmA_Trx), co-expressed with the His-tagged B1 protein (6xHis_PsmB1) and PsmB2 in *E. coli*, and co-purified with 6xHis_PsmB1 tightly bound to the leader part of PsmA.^5^ In the presence of the wild-type B2 protein, most of MBP_PsmA_Trx was cleaved forming MBP_PsmA-Leader. Truncation of the N-terminal α-helix (PsmB2-ΔN14) did not affect the activity. For hydrophilic mutations, we chose one representative residue each from the predicted protein surface, the hydrophobic core, and the potential interaction patch with the B1 protein (Psm B2 Ile33, Val72, and V142, respectively). The I33K surface mutant (corresponding to TfuB2 L31K) retained a comparable activity to that of the wild-type protein. In contrast, the V72K mutant (TfuB2 V66K) in the hydrophobic core and the V142K mutant (TfuB2 L134K) on the potential interaction patch completely lost the peptidase activity. These results are highly consistent with the structure prediction and the results obtained with TfuB2, indicating that the structural features, most notable, the interaction patch, are conserved among the B2 proteins from different organisms.

**Figure 4.**
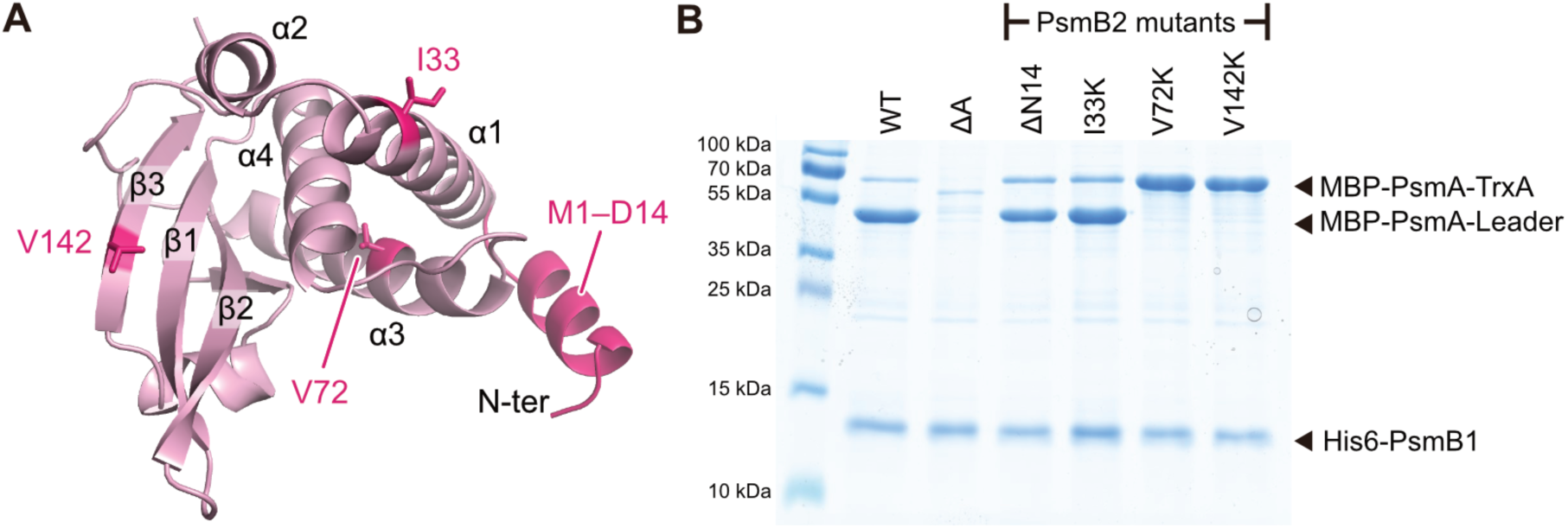
Structure prediction and mutational analysis of PsmB2. (A) The structural model of PsmB2 predicted by AlphaFold2. (B) The *in vivo* activity analysis of PsmB2 mutants. The reaction in lane ΔA was performed without PsmA.

## CONCLUSION

In this study, we have performed a quick semi-comprehensive mutant analysis of TfuB2, by combining three technologies: assembly PCR, cell-free protein expression, and determination of the enzymatic activity without purification. The results confirmed the structure prediction performed by AlphaFold2 and identified a possible interaction surface with TfuB1. Although such a rapid cell-free analysis obviously would not work with all proteins, the combination of the precise structure prediction and the cell-free semi-comprehensive mutant analysis is a powerful approach that would accelerate the biochemical research of a large variety of proteins providing informative results in a matter of days (Day 1: AlphaFold2 prediction and primer design; Day 2: gene assembly, cell-free expression, and functional analysis).

## Supporting Information

*Supporting Information Available*: This material is available free of charge *via* the Internet.

## Acknowledgements

S.T. was supported by the Inamori Foundation and the Koyanagi Foundation.

