## Supplementary information for "Cell-free mutant analysis combined with structure prediction of a lasso peptide biosynthetic protein B2"

### METHODS

#### TfuA cleavage assay in the PURE system

The primers and templates used to amplify the genes of TfuA-sfGFP, TfuB1, and TfuB2 variants are summarized in Supplementary Tables 1 and 2. The sequences of TfuA, TfuB1, and TfuB2 were optimized for protein expression in *E. coli* by the supplier's website (Thermo Fisher Scientific). The constructs of TfuA-sfGFP, TfuB1, and wild-type/truncated TfuB2 were amplified from the synthetic genes by standard PCR reactions (Supplementary Table 3). The TfuB2 constructs with amino acid substitutions were prepared by assembly PCR[1] from fragmented oligo DNAs (Supplementary Table 1–3). A T7 promoter and a ribosome binding site were also added during the PCR reactions. All constructs have an N-terminal extension encoded by a 30 nt A-rich sequence for stable protein expression in the PURE system.

For cell-free analysis of the TfuB2 activity, 1.25 µL solution I, 0.125 µL solution II, and 0.25 µL solution III of PUREfrex 2.0 (GeneFrontier) were mixed with 0.125 µL RNase inhibitor (Nacalai Tesque, Japan), 0.1 µL each of templates of TfuA-sfGFP, TfuB1, and TfuB2, and 0.45 µL $H_2O$. The samples were incubated at 37 ºC for 1 hour, and at 50 ºC for 1 hour to complete the expression and peptidase reaction, respectively. A mixture of 2.5 µL $H_2O$ and 2.5 µL 4 x LDS sample buffer (Thermo Fisher Scientific) was added to the incubated sample. 2 µL of each sample was loaded to SDS-PAGE gels (NuPAGE 4–12% Bis-Tris gel, Thermo Fisher Scientific) without heating to avoid denaturation of sfGFP. The gels were run at 100 V for 60 min and analyzed by Amersham Typhoon (GE Healthcare). The analyses for truncation variants and lysine substitutions of TfuB2 were performed before the release of AlphaFold2, except for reproduction of the results for publication. The alanine substitutions of the active site residues were designed after examining the predicted structure of TfuB2.

**Expression and solubility test of lysine mutants of TfuB2 in the PURE system**

To express fluorescently labeled proteins, we added FluoroTect GreenLys, tRNA charged with a fluorescently-labeled lysine (Promega), to the PURE system. 2.5 μL solution I, 0.25 μL solution II, 0.5 μL solution III of PUREfrex 2.0 were mixed with 0.25 μL RNase inhibitor, 0.2 μL FluoroTect GreenLys, and 1.3 μL templates for the TfuB2 variants. The samples were incubated at 37 ºC for 1 hour to complete protein expression. Then, they were mixed with 1.0 μL 1 mg/mL RNase A and incubated at 37 ºC for 30 min to digest the extra amount of FluoroTect GreenLys. For the solubility test, the samples were further incubated at 50 ºC for 15 min to denature the proteins from PUREfrex and divided into supernatants and precipitations by centrifugation. A mixture of 3 μL $H_2O$ and 3 μL 4 x LDS sample buffer was added to each sample. The whole amounts of the samples were loaded to SDS-PAGE gels (NuPAGE 4–12% Bis-Tris gel) after heating at 95 ºC for 5 min. The gels were run at 180 V for 30 min and analyzed by Amersham Typhoon (GE Healthcare).

**Structure predictions**

AlphaFold2 was employed to predict the three-dimensional structures of TfuB2, PsmB2, and TfuA-Leader•TfuB1•TfuB2 protein-protein complex.[2,3] AlphaFold2 source code was retrieved from its github repository (https://github.com/deepmind/alphafold) and locally installed on a Linux workstation with NVIDIA Quadro RTX8000 GPU. Structure prediction was performed using "*full_dbs*" as db_preset parameter. TfuB2 and PsmB2 proteins were predicted using "*monomer*" mode of AlphaFold2, while "*multimer*" mode was employed for the prediction of TfuA-Leader•TfuB1•TfuB2 protein-protein complex. For all three protein targets, five models were generated and the model with the best pLDDT score, a confidence measure from AlphaFold2, was selected. pLDDT scores were extracted using "plddt2csv.py" python script from https://github.com/CYP152N1/plddt2csv.

***In vivo* cleavage assay of PmsA**

Sequences of the oligonucleotides used in *in vivo* mutation study of PsmB2 are listed in Supplemental Table 4. The investigation of leader peptide cleavage was performed using pAIR plasmid[4]. Mutations in *psmB2* gene were introduced by overlap extension PCR amplification[5] with the pAIR vector encoding wild-type *psmB2* gene as a template. PCR product then was digested with BamHI and XhoI FastDigest restriction endonucleases (Thermo Scientific) and inserted into pAIR vector which was linearized using the same restriction enzymes.

*E. coli* BL21(DE3) cells harboring the pAIR vector were grown at 37°C with 180 rpm shaking in 15 mL of 2xYT medium supplemented with 50 mg/mL of kanamycin until $OD_{600}$ reached 0.6. Then protein expression was induced by addition of 1 mM of isopropyl-β-thiogalactopyranoside, followed by growth at 37 °C with 180 rpm shaking for 2.5 hours. Cells were harvested by centrifugation, resuspended in 1 mL of lysis buffer (20 mM Tris-HCl, pH 8.0; 150 mM NaCl; 5 mM imidazole), and lysed by sonication. Cleared cell lysates were obtained by centrifugation, mixed with 50 µL of TALON Superflow Metal Affinity Resin (Takara) pre-equilibrated with lysis buffer, and incubated at 4°C with gentle rotation for 45 min. The resin then was washed with lysis buffer and the protein complexes were eluted with 100 µL of elution buffer (500 mM Imidazole, pH 8.0; 20 mM Tris-HCl, pH 8.0; 150 mM NaCl). Eluted fractions were analyzed with 13% SDS PAGE. The gels were stained with Instant*Blue* gel stain (ISBT) according to the manufacturer's protocol.

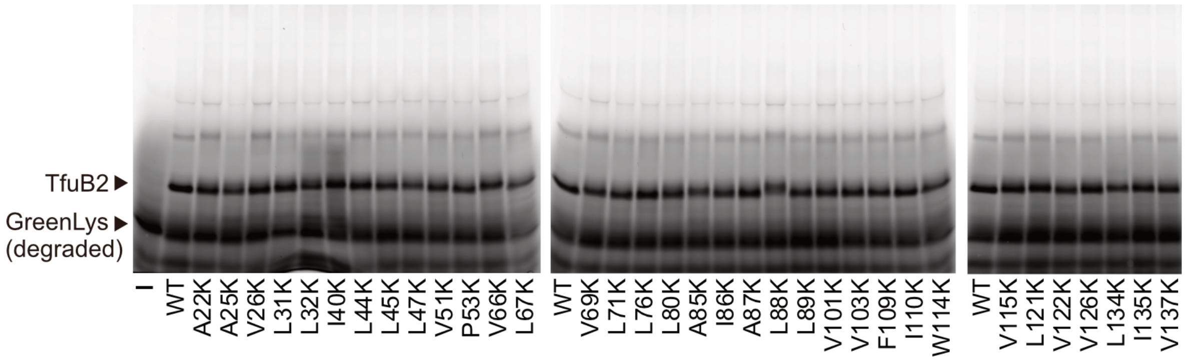


**Supplementary Figure 1. Expression of TfuB2 mutants the PURE system.** The expressions of TfuB2-ΔN4 and its hydrophilic mutants in the PURE system were monitored using an *in vitro* translation labeling system, FluoroTect GreenLys (Promega), which contains tRNA charged with a fluorescently-labeled lysine.

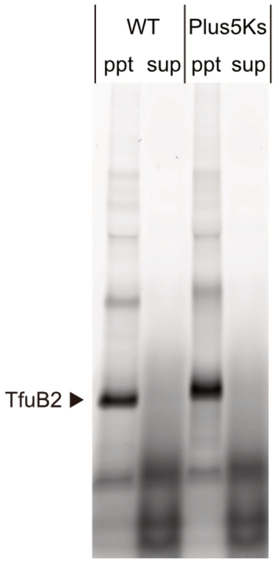


**Supplementary Figure 2. Solubility of TfuB2 in the PURE system.** Solubilities of TfuB2-ΔN4 and a hyper-hydrophilic variant (Plus5Ks) were investigated by centrifugation after incubation at 50ºC (the temperature for the activity test of TfuB2) for 15 min. Both the wild-type TfuB2 and the Plus5Ks mutant completely precipitated.

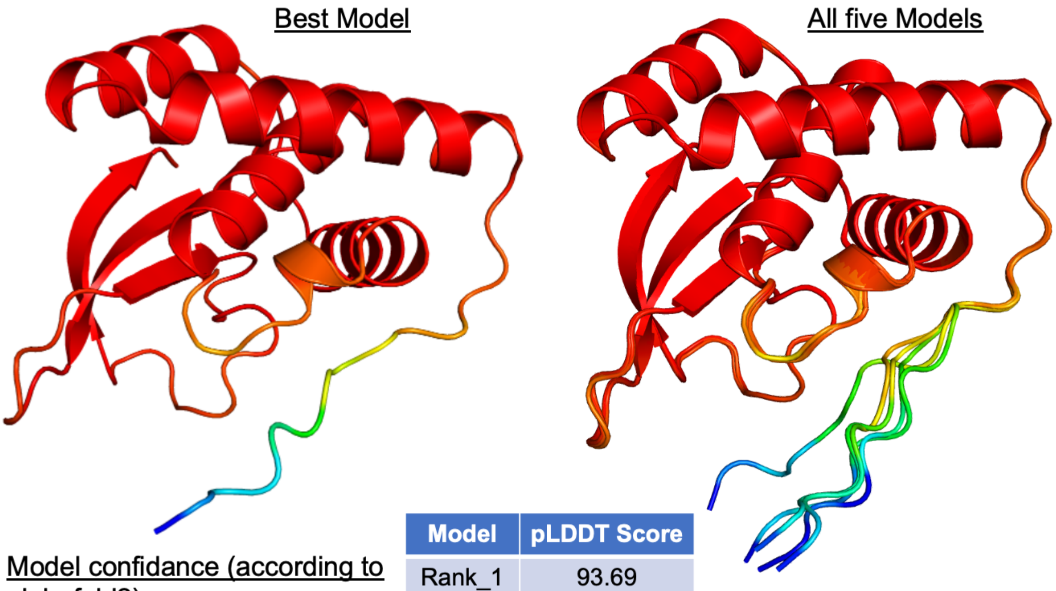


Model confidance (according to alphafold2)

Very high (pLDDTS > 90)
Confidant (90 > pLDDTS > 70)
Low (70 > pLDDTS > 50)
Very low (pLDDTS < 50)

| Model | pLDDT Score |
|---|---|
| Rank_1 | 93.69 |
| Rank_2 | 93.68 |
| Rank_3 | 93.01 |
| Rank_4 | 91.93 |
| Rank_5 | 91.10 |

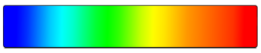


**Supplementary Figure 3. AlphaFold2 predicted structure of TfuB2.** Models are colored according to pLDDT score and average pLDDT score for the five predicted models is given in the table at the bottom center.

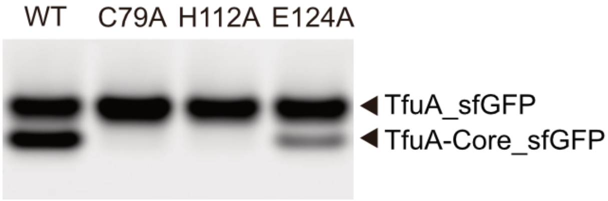


**Supplementary Figure 4. Activity test of the catalytic site mutants in the PURE system.** The TfuA cleavage activities of the catalytic site mutants of TfuB2 were monitored by cell-free co-expression with TfuA-sfGFP and TfuB1.

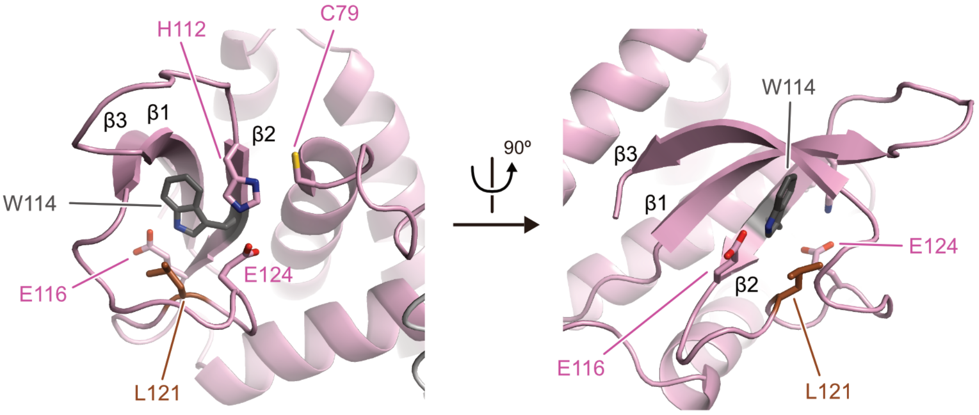


**Supplementary Figure 5. Conformation of the loop between β2 and β3.** Close-up view of the loop between β2 and β3 (117–130) in the predicted TfuB2 structure. Trp114 and L121 contact each other and form a small core-like part behind the loop. The E116A mutant reportedly lost the activity[6] despite the fact that it does not directly contact the active site in the predicted structure. Instead, Glu116 contacts Trp114, likely fixing the position/direction of the hydrophobic residue, although the accuracy of the prediction in such detail/resolution has not been assured.

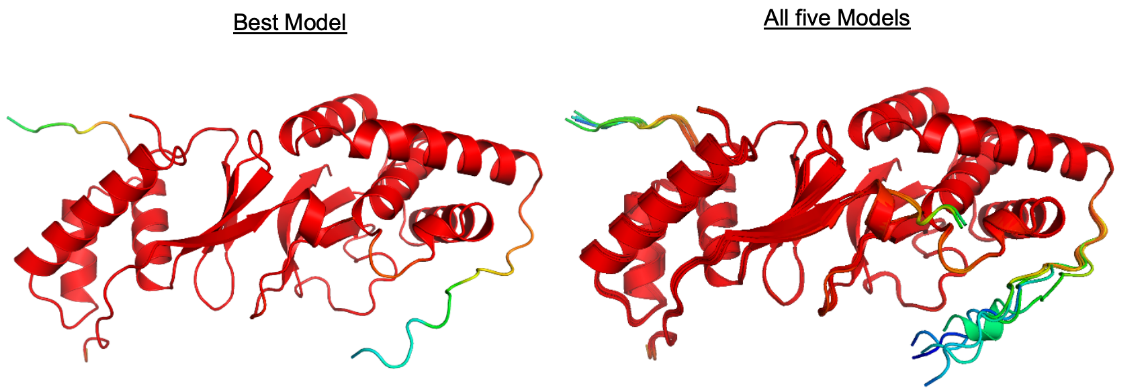


Model confidance (according to alphafold2)

Very high (pLDDTS > 90)
Confidant (90 > pLDDTS > 70)
Low  (70 > pLDDTS > 50)
Very low (pLDDTS < 50)

| Model | pLDDT Score |
|---|---|
| Rank_1 | 0.88 |
| Rank_2 | 0.87 |
| Rank_3 | 0.87 |
| Rank_4 | 0.87 |
| Rank_5 | 0.86 |

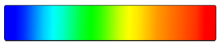


**Supplementary Figure 6. AlphaFold2 predicted structure of TfuA-Leader•TfuB1•TfuB2 complex.** Models are colored according to pLDDT score and average pLDDT score for the five predicted models is given in the table at the bottom center.

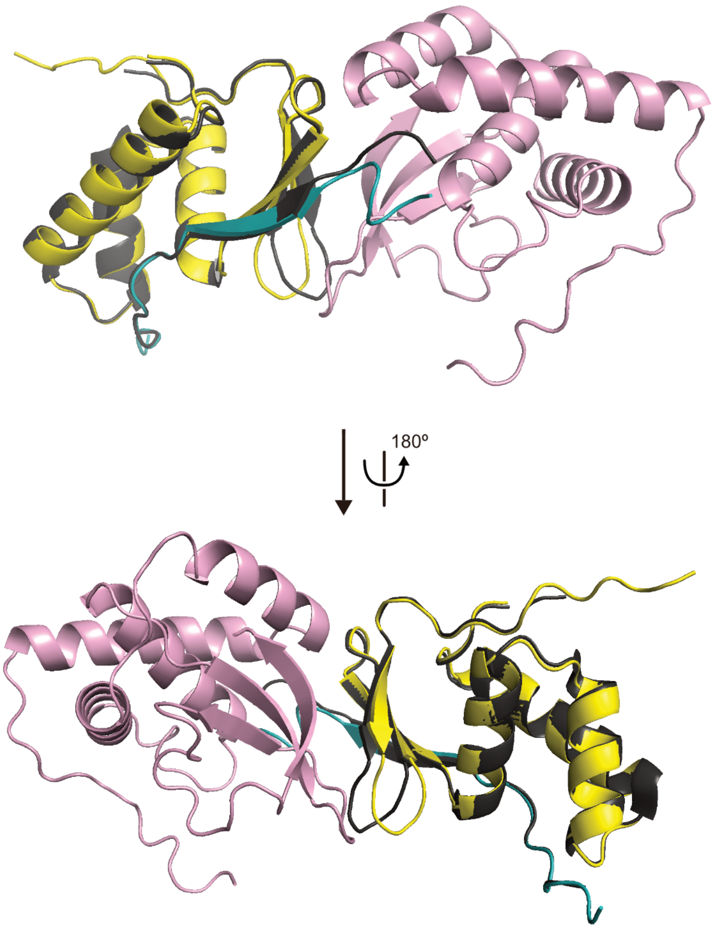


**Supplementary Figure 7. Superimposition of the predicted structure of TfuA-Leader•TfuB1•TfuB2 and the crystal structure of TfuA-Leader•TfuB1.** The previously reported crystal structure of the TfuA-Leader•TfuB1 complex (black, PDB: 6JX3)[4] is superimposed with the structural model of TfuA-Leader•TfuB1•TfuB2 predicted by AlphaFold2. The predicted structure is colored in the same way as in Figure 3 (TfuA-Leader: cyan; TfuB1: yellow; TfuB2: pink).

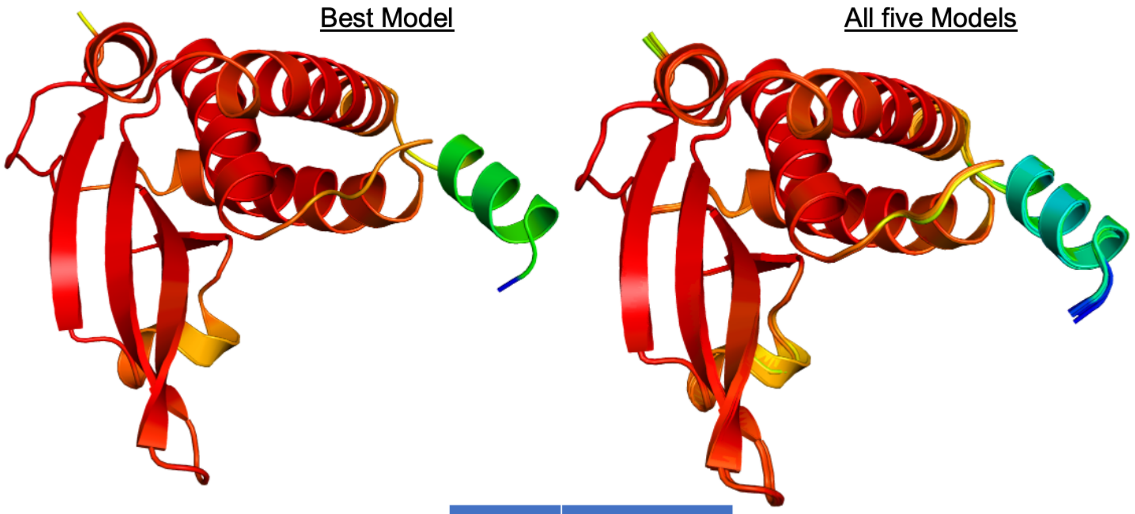


Model confidance (according to alphafold2)

Very high (pLDDTS > 90)
Confidant (90 > pLDDTS > 70)
Low (70 > pLDDTS > 50)
Very low (pLDDTS < 50)

| Model | pLDDT Score |
|---|---|
| Rank_1 | 93.06 |
| Rank_2 | 91.91 |
| Rank_3 | 91.91 |
| Rank_4 | 91.86 |
| Rank_5 | 91.07 |

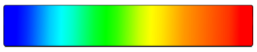


**Supplementary Figure 8. AlphaFold2 predicted structure of PsmB2.** Models are colored according to pLDDT score and average pLDDT score for the five predicted models is given in the Table.

Supplementary Table 1. Synthetic genes

| Gene | Manufacturer | Sequence |
|---|---|---|
| TfuA-sfGFP-gene | Thermo Fisher | GCGAATTAATACGACTCACTATAGGGCTTAAGTATAAGGA GGAAAAAATATGAAAATAAAAACAGGAGCACGCAATAAC ATGGAAAAGAAGAAATATACCGCACCGCAGCTGGCAAAA GTTGGTGAATTTAAAGAAGCCACCGGTTGGTATACCGCAG AATGGGGTTTAGAACTGATTTTTGTTTTTCCGCGTTTTATT CATCACCATCATCACCACTCCGCGGCTCTTGAAGTCCTCTT TCAGGGACCCATGAGCAAAGGAGAAGAACTTTTCACTGGA GTTGTCCCAATTCTTGTTGAATTAGATGGTGATGTTAATGG GCACAAATTTTCTGTCCGTGGAGAGGGTGAAGGTGATGCT ACAAACGGAAAACTCACCCTTAAATTTATTTGCACTACTGG AAAACTACCTGTTCCATGGCCAACACTTGTCACTACTCTGA CCTATGGTGTTCAATGCTTTTCCCGTTATCCGGATCACATG AAACGGCATGACTTTTTCAAGAGTGCCATGCCCGAAGGTT ATGTACAGGAACGCACTATATCTTTCAAAGATGACGGGAC CTACAAGACGCGTGCTGAAGTCAAGTTTGAAGGTGATACC CTTGTTAATCGTATCGAGTTAAAAGGTATTGATTTTAAAGA AGATGGAAACATTCTCGGACACAAACTCGAGTACAACTTT AACTCACACAATGTATACATCACGGCAGACAAACAAAAGA ATGGAATCAAAGCTAACTTCAAAATTCGCCACAACGTTGA AGATGGTTCCGTTCAACTAGCAGACCATTATCAACAAAAT ACTCCAATTGGCGATGGCCCTGTCCTTTTACCAGACAACCA TTACCTGTCGACACAATCTGTCCTTTCGAAAGATCCCAACG AAAAGCGTGACCACATGGTCCTTCTTGAGTTTGTAACTGCT GCTGGGATTACACATGGCATGGATGAGCTCTACAAAGGTA GCTAATAAggaccgaattctgtacaggcc<br>(T7 promoter–SD–A-rich sequence–linker–TfuA–linker–sfGFP) |
| TfuB1-gene<br>(1–90, M1V) | Thermo Fisher | GTTGAAACCACCGGTGCAGAATTTCGTCTGCGTCCGGAAAT TAGCGTTGCACAGACCGATTATGGTATGGTTCTGCTGGATG GTCGTAGCGGTGAATATTGGCAGCTGAATGATACCGCAGC ACTGATTGTTCAGCGTCTGCTGGACGGCCATAGTCCGGCAG ATGTTGCACAGTTTCTGACCAGCGAATATGAAGTTGAACGT ACCGATGCAGAACGTGATATTGCAGCACTGGTTACCAGCC TGAAAGAAAATGGTATGGCACTGCCG<br>(The original start codon of TfuB1 was GTG and it was converted to GTT in the codon optimization process by the supplier's website, which caused a mutation to valine without any significant effect)[4] |
| TfuB2-gene<br>(5–138) | Thermo Fisher | GTTGTTCTGCAGCGTAGCAATGTTCGTCTGAGCTGGCGTAC CAAATGGGCAGCACGTTGTGCAGTTGGTGCAGCACGTCTG CTGGCACGTAAACCGCCTGAACGTATTCGTGCAACCCTGCT GCGTCTGCGTGGTGAAGTTCGTCCGGCAACCTATGAAGAG GCAAAAGCAGCACGTGATGCCGTTCTGGCAGTTAGCCTGC GTTGTGCCGGTCTGCGTGCATGTCTGCAGCGCAGCCTGGCA ATTGCACTGCTGTGTCGTATGCGTGGCACCTGGGCAACCTG GTGTGTTGGTGTTCCGCGTCGTCCGCCTTTTATTGGTCATGC ATGGGTTGAAGCAGAAGGTCGTCTGGTTGAAGAGGGTGTT GGTTATGATTATTTCAGCCGTCTGATTACCGTGGAT<br>(Overlap regions between the fragmented oligo DNAs are colored differently. Also see Supplementary Table 2 and 3) |

Supplementary Table 2. DNA oligos for cell-free experiments

| Name | Sequence |
|---|---|
| PURE-F | GCGAATTAATACGACTCACTATAGGGCTTAAGTATAAGGAGGAAAAAATATGAAAATAAAAACAGGAGCACGCAATAAC (T7 promoter–SD–A-rich sequence) |
| PURE-R | ggcctgtacagaattcggtccTTATTA |
| TfuB1-F | ATGAAAATAAAAACAGGAGCACGCAATAACGTTGAAACCACCGGTGCAGAATTTC |
| TfuB1-R | ggcctgtacagaattcggtccTTATTACGGCAGTGCCATACCATTTTCTTTC |
| TfuB2-F | ATGAAAATAAAAACAGGAGCACGCAATAACatgtctgaaaatGTTGTTCTGCAGCGTAGCAATGTTCG |
| TfuB2-R | ggcctgtacagaattcggtccTTATTAATCCACGGTAATCAGACGGCTG |
| TfuB2-ΔN4-F | ATGAAAATAAAAACAGGAGCACGCAATAACGTTGTTCTGCAGCGTAGCAATGTTC |
| TfuB2-ΔN12-F | ATGAAAATAAAAACAGGAGCACGCAATAACCTGAGCTGGCGTACCAAATGGG |
| TfuB2-ΔN18-F | ATGAAAATAAAAACAGGAGCACGCAATAACAAATGGGCAGCACGTTGTGCA |
| TfuB2-ΔN24-F | ATGAAAATAAAAACAGGAGCACGCAATAACGCAGTTGGTGCAGCACGTC |
| TfuB2-ΔN30-F | ATGAAAATAAAAACAGGAGCACGCAATAACCTGCTGGCACGTAAACCGC |
| TfuB2-ΔC5-R | ggcctgtacagaattcggtccTTATTAACGGCTGAAATAATCATAACCAACACCCTC |
| TfuB2-ΔC14-R | ggcctgtacagaattcggtccTTATTATCTTCAACCAGACGACCTTCTGCTTC |
| B2frag1 (Forward) | ATGAAAATAAAAACAGGAGCACGCAATAACGTTGTTCTGCAGCGTAGCAATGTTCGTCTGAGCTGGCGTA |
| B2frag2 (Forward) | AATGTTCGTCTGAGCTGGCGTACCAAATGGGCAGCACGTTGTGCAGTTGGTGCAGCACGTCTGCTGGCAC |
| B2frag3 (Forward) | CAGCACGTCTGCTGGCACGTAAACCGCCTGAACGTATTCGTGCAACCCTGCTGCGTCTGCGTGGTGAAGT |
| B2frag4 (Forward) | TGCGTCTGCGTGGTGAAGTTCGTCCGGCAACCTATGAAGAGGCAAAAGCAGCACGTGATGCCGTTCTGGC |
| B2frag5 (Reverse) | CAGGCTGCGCTGCAGACATGCACGCAGACCGGCACAACGCAGGCTAACTGCCAGAACGGCATCACGTG |
| B2frag6 (Reverse) | CCAACACACCAGGTTGCCCAGGTGCCACGCATACGACACAGCAGTGCAATTGCCAGGCTGCGCTGCAGAC |
| B2frag7 (Reverse) | CTTCTGCTTCAACCCATGCATGACCAATAAAGGCGGACGACGCGGAACACCAACACACCAGGTTGCCCA |
| B2frag8 (Reverse) | GACGGCTGAAATAATCATAACCAACACCCTCTTCAACCAGACGACCTTCTGCTTCAACCCATGCATGACC |
| B2frag9 (Reverse) | ggcctgtacagaattcggtccTTATTAATCCACGGTAATCAGACGGCTGAAATAATCATAACCAACACCC |
| B2frag2-A22K | AATGTTCGTCTGAGCTGGCGTACCAAATGGGCAaaaCGTTGTGCAGTTGGTGCAGCACGTCTGCTGGCAC |
| B2frag2-A25K | AATGTTCGTCTGAGCTGGCGTACCAAATGGGCAGCACGTTGTaaaGTTGGTGCAGCACGTCTGCTGGCAC |
| B2frag2-V26K | AATGTTCGTCTGAGCTGGCGTACCAAATGGGCAGCACGTTGTGCAaaaGGTGCAGCACGTCTGCTGGCAC |
| B2frag2-L31K | AATGTTCGTCTGAGCTGGCGTACCAAATGGGCAGCACGTTGTGCAGTTGGTGCAGCACGTaaaCTGGCAC |
| B2frag3-L31K | CAGCACGTaaaCTGGCACGTAAACCGCCTGAACGTATTCGTGCAACCCTGCTGCGTCTGCGTGGTGAAGT |
| B2frag2-L32K | AATGTTCGTCTGAGCTGGCGTACCAAATGGGCAGCACGTTGTGCAGTTGGTGCAGCACGTCTGaaaGCAC |
| B2frag3-L32K | CAGCACGTCTGaaaGCACGTAAACCGCCTGAACGTATTCGTGCAACCCTGCTGCGTCTGCGTGGTGAAGT |
| B2frag3-I40K | CAGCACGTCTGCTGGCACGTAAACCGCCTGAACGTaaaCGTGCAACCCTGCTGCGTCTGCGTGGTGAAGT |

| | |
|---|---|
| B2frag3-L44K | CAGCACGTCTGCTGGCACGTAAACCGCCTGAACGTATTCGTGCAACCaaaCTGCGTCTGCGTGGTGAAGT |
| B2frag3-L45K | CAGCACGTCTGCTGGCACGTAAACCGCCTGAACGTATTCGTGCAACCCTGaaaCGTCTGCGTGGTGAAGT |
| B2frag4-L45K | aaCGTCTGCGTGGTGAAGTTCGTCCGGCAACCTATGAAGAGGCAAAAGCAGCACGTGATGCCGTTCTGGC |
| B2frag3-L47K | CAGCACGTCTGCTGGCACGTAAACCGCCTGAACGTATTCGTGCAACCCTGCTGCGTaaaCGTGGTGAAGT |
| B2frag4-L47K | TGCGTaaaCGTGGTGAAGTTCGTCCGGCAACCTATGAAGAGGCAAAAGCAGCACGTGATGCCGTTCTGGC |
| B2frag3-V51K | CAGCACGTCTGCTGGCACGTAAACCGCCTGAACGTATTCGTGCAACCCTGCTGCGTCTGCGTGGTGAAaa |
| B2frag4-V51K | TGCGTCTGCGTGGTGAAaaaCGTCCGGCAACCTATGAAGAGGCAAAAGCAGCACGTGATGCCGTTCTGGC |
| B2frag4-P53K | TGCGTCTGCGTGGTGAAGTTCGTaaaGCAACCTATGAAGAGGCAAAAGCAGCACGTGATGCCGTTCTGGC |
| B2frag4-V66K | TGCGTCTGCGTGGTGAAGTTCGTCCGGCAACCTATGAAGAGGCAAAAGCAGCACGTGATGCCaaaCTGGC |
| B2frag5-V66K | CAGGCTGCGCTGCAGACATGCACGCAGACCGGCACAACGCAGGCTAACTGCCAGtttGGCATCACGTG |
| B2frag4-L67K | TGCGTCTGCGTGGTGAAGTTCGTCCGGCAACCTATGAAGAGGCAAAAGCAGCACGTGATGCCGTTaaaGC |
| B2frag5-L67K | CAGGCTGCGCTGCAGACATGCACGCAGACCGGCACAACGCAGGCTAACTGCtttAACGGCATCACGTG |
| B2frag5-V69K | CAGGCTGCGCTGCAGACATGCACGCAGACCGGCACAACGCAGGCTtttTGCCAGAACGGCATCACGTG |
| B2frag5-L71K | CAGGCTGCGCTGCAGACATGCACGCAGACCGGCACAACGtttGCTAACTGCCAGAACGGCATCACGTG |
| B2frag5-L76K | CAGGCTGCGCTGCAGACATGCACGtttACCGGCACAACGCAGGCTAACTGCCAGAACGGCATCACGTG |
| B2frag5-L80K | CAGGCTGCGCTGtttACATGCACGCAGACCGGCACAACGCAGGCTAACTGCCAGAACGGCATCACGTG |
| B2frag6-L80K | CCAACACACCAGGTTGCCCAGGTGCCACGCATACGACACAGCAGTGCAATTGCCAGGCTGCGCTGtttAC |
| B2frag6-A85K | CCAACACACCAGGTTGCCCAGGTGCCACGCATACGACACAGCAGTGCAATtttCAGGCTGCGCTGCAGAC |
| B2frag6-I86K | CCAACACACCAGGTTGCCCAGGTGCCACGCATACGACACAGCAGTGCtttTGCCAGGCTGCGCTGCAGAC |
| B2frag6-A87K | CCAACACACCAGGTTGCCCAGGTGCCACGCATACGACACAGCAGtttAATTGCCAGGCTGCGCTGCAGAC |
| B2frag6-L88K | CCAACACACCAGGTTGCCCAGGTGCCACGCATACGACACAGtttTGCAATTGCCAGGCTGCGCTGCAGAC |
| B2frag6-L89K | CCAACACACCAGGTTGCCCAGGTGCCACGCATACGACAtttCAGTGCAATTGCCAGGCTGCGCTGCAGAC |
| B2frag6-V101K | CCtttACACCAGGTTGCCCAGGTGCCACGCATACGACACAGCAGTGCAATTGCCAGGCTGCGCTGCAGAC |
| B2frag7-V101K | CTTCTGCTTCAACCCATGCATGACCAATAAAAGGCGGACGACGCGGAACACCtttACACCAGGTTGCCCA |
| B2frag7-V103K | CTTCTGCTTCAACCCATGCATGACCAATAAAAGGCGGACGACGCGGtttACCAACACACCAGGTTGCCCA |
| B2frag7-F109K | CTTCTGCTTCAACCCATGCATGACCAATtttAGGCGGACGACGCGGAACACCAACACACCAGGTTGCCCA |
| B2frag7-I110K | CTTCTGCTTCAACCCATGCATGACCtttAAAAGGCGGACGACGCGGAACACCAACACACCAGGTTGCCCA |
| B2frag7-W114K | CTTCTGCTTCAACtttTGCATGACCAATAAAAGGCGGACGACGCGGAACACCAACACACCAGGTTGCCCA |
| B2frag8-W114K | GACGGCTGAAATAATCATAACCAACACCCTCTTCAACCAGACGACCTTCTGCTTCAACtttTGCATGACC |

| | |
|---|---|
| B2frag7-V115K | CTTCTGCTTCtttCCATGCATGACCAATAAAAGGCGGACGACGCGGAACACC AACACACCAGGTTGCCCA |
| B2frag8-V115K | GACGGCTGAAATAATCATAACCAACACCCTCTTCAACCAGACGACCTTCTG CTTCtttCCATGCATGACC |
| B2frag8-L121K | GACGGCTGAAATAATCATAACCAACACCCTCTTCAACtttACGACCTTCTGCT TCAACCCATGCATGACC |
| B2frag8-V122K | GACGGCTGAAATAATCATAACCAACACCCTCTTCtttCAGACGACCTTCTGCT TCAACCCATGCATGACC |
| B2frag8-V126K | GACGGCTGAAATAATCATAACCtttACCCTCTTCAACCAGACGACCTTCTGCT TCAACCCATGCATGACC |
| B2frag9-V126K | ggcctgtacagaattcggtccTTATTAATCCACGGTAATCAGACGGCTGAAATAATCAT AACCtttACCC |
| B2frag8-L134K | tACGGCTGAAATAATCATAACCAACACCCTCTTCAACCAGACGACCTTCTGC TTCAACCCATGCATGACC |
| B2frag9-L134K | ggcctgtacagaattcggtccTTATTAATCCACGGTAATtttACGGCTGAAATAATCATAA CCAACACCC |
| B2frag9-I135K | ggcctgtacagaattcggtccTTATTAATCCACGGTtttCAGACGGCTGAAATAATCATAA CCAACACCC |
| B2frag9-V137K | ggcctgtacagaattcggtccTTATTAATCtttGGTAATCAGACGGCTGAAATAATCATAA CCAACACCC |
| B2frag6-C79A | CCAACACACCAGGTTGCCCAGGTGCCACGCATACGACACAGCAGTGCAATT GCCAGGCTGCGCTGCAGAg |
| B2frag7-H112A | CTTCTGCTTCAACCCATGCAgcACCAATAAAAGGCGGACGACGCGGAACAC CAACACACCAGGTTGCCCA |
| B2frag8-H112A | GACGGCTGAAATAATCATAACCAACACCCTCTTCAACCAGACGACCTTCTG CTTCAACCCATGCAgcACC |
| B2frag8-E124A | GACGGCTGAAATAATCATAACCAACACCCgCTTCAACCAGACGACCTTCTG CTTCAACCCATGCATGACC |

Supplementary Table 3. PCR schemes for TfuA-sfGFP, TfuB1, and TfuB2 variants

| Gene | Round | Primer (Forward) (× 1) | Template or Fragments (× 0.1) | Primer (Reverse) (× 1) |
|---|---|---|---|---|
| TfuA-sfGFP | | PURE-F | TfuA-sfGFP-gene | PURE-R |
| TfuB1 | PCR1 | TfuB1-F | TfuB1-gene | TfuB1-R |
| | PCR2 | PURE-F | PCR1 | PURE-R |
| TfuB2 | PCR1 | TfuB2-F | TfuB2-gene | TfuB2-R |
| | PCR2 | PURE-F | PCR1 | PURE-R |
| TfuB2-ΔN4 | PCR1 | TfuB2-ΔN4-F | TfuB2-gene | TfuB2-R |
| | PCR2 | PURE-F | PCR1 | PURE-R |
| TfuB2-ΔN12 | PCR1 | TfuB2-ΔN12-F | TfuB2-gene | TfuB2-R |
| | PCR2 | PURE-F | PCR1 | PURE-R |
| TfuB2-ΔN18 | PCR1 | TfuB2-ΔN18-F | TfuB2-gene | TfuB2-R |
| | PCR2 | PURE-F | PCR1 | PURE-R |
| TfuB2-ΔN24 | PCR1 | TfuB2-ΔN24-F | TfuB2-gene | TfuB2-R |
| | PCR2 | PURE-F | PCR1 | PURE-R |
| TfuB2-ΔN30 | PCR1 | TfuB2-ΔN30-F | TfuB2-gene | TfuB2-R |
| | PCR2 | PURE-F | PCR1 | PURE-R |
| TfuB2-ΔC5 | PCR1 | TfuB2-F | TfuB2-gene | TfuB2-ΔC5-R |
| | PCR2 | PURE-F | PCR1 | PURE-R |
| TfuB2-ΔC14 | PCR1 | TfuB2-F | TfuB2-gene | TfuB2-ΔC14-R |
| | PCR2 | PURE-F | PCR1 | PURE-R |
| Lysine/Alanine substitutions of TfuB2 | | PURE-F | B2frag1–8 with corresponding mutations | B2frag9 with corresponding mutations |

Supplementary Table 4. Primers used for *in vivo* mutation analysis of PsmB2

| Primer name | Sequence |
|---|---|
| PsmB2_trunc_F | ttaGGAGTAGGATCGATGGGAACAAAATTA |
| PsmB2_trunc_R | TAATTTTGTTCCCATCGATCCTACTCCTAA |
| PsmB2_I33K_F | TTAGGCTGGGCTCGTaaaCTTAAAAGTATACCG |
| PsmB2_I33K_R | CGGTATACTTTTAAGtttACGAGCCCAGCCTAA |
| PsmB2_V72K_F | AAAATTTCTCAAGCAaaaCATATGATGAGCCGT |
| PsmB2_V72K_R | ACGGCTCATCATATGtttTGCTTGAGAAATTTT |
| PsmB2_V142K_F | AAGAGATTTACTGTTaaaGGTAAATTTGCAAAA |
| PsmB2_V142K_R | TTTTGCAAATTTACCtttAACAGTAAATCTCTT |
| T7Duet2F | TTGTACACGGCCGCATAAT |
| T7ter | GCTAGTTATTGCTCAGCGG |
